# The transcriptional and translational landscape of HCoV-OC43 infection

**DOI:** 10.1101/2024.01.20.576440

**Authors:** Stefan Bresson, Emanuela Sani, Alicja Armatowska, Charles Dixon, David Tollervey

## Abstract

The coronavirus HCoV-OC43 circulates continuously in the human population and is a frequent cause of the common cold. Here, we generated a high-resolution atlas of the transcriptional and translational landscape of OC43 during a time course following infection of human lung fibroblasts. Using ribosome profiling, we quantified the relative expression of the canonical open reading frames (ORFs) and identified previously unannotated ORFs. These included several short upstream ORFs and a putative ORF nested inside the M gene. In parallel, we analyzed the cellular response to infection. Endoplasmic reticulum (ER) stress response genes were transcriptionally and translationally induced beginning 12 and 18 hours post infection, respectively. By contrast, conventional antiviral genes mostly remained quiescent. At the same time points, we observed accumulation and increased translation of noncoding transcripts normally targeted by nonsense mediated decay (NMD), suggesting NMD is suppressed during the course of infection. This work provides resources for deeper understanding of OC43 gene expression and the cellular responses during infection.

## INTRODUCTION

To date, seven coronaviruses have been found to infect humans and cause respiratory disease, ranging from mild cold to severe pneumonia. These seven coronaviruses are broadly grouped into two distinct lineages: the alphacoronaviruses (HCoV-229E and HCoV-NL63) and betacoronaviruses (HCoV-OC43, HCoV-HKU1, SARS-CoV, MERS-CoV, and SARS-CoV-2). Each virus has a distinct complement of accessory, and sometimes structural, genes and they frequently use different receptors for entry^1^. Despite these differences, the core replication mechanisms are highly conserved^2^. During early infection, the incoming, positive-strand genomic RNA (+)gRNA is the only viral genetic material (Fig. S1a). The (+)gRNA is initially translated to generate the large polyproteins ORF1A and ORF1AB, the latter regulated by a -1 frameshift element that allows ribosomes to continue translating past the stop codon at the end of ORF1A. The polyproteins are cleaved by two viral proteases to yield a set of “non-structural proteins” (NSP1 through NSP16), including the RNA processing enzymes and components of the viral RNA-dependent RNA polymerase (RdRp). The viral RdRp uses the (+)gRNA to synthesize a negative-strand intermediate, (-)gRNA, which in turn serves as a template for synthesis of additional (+)gRNAs. In addition, RdRp synthesizes shorter negative-strand subgenomic RNAs, (-)sgRNAs, by prematurely disengaging from the template strand before resuming transcription at a downstream site (Fig. S1b). This process, known as discontinuous transcription, is guided by the presence of transcription regulatory sequences (TRSs). The resulting (-)sgRNAs serve as templates for synthesis of plus-strand subgenomic RNAs, (+)sgRNAs. These act as monocistronic mRNAs for translation of one or more accessory proteins and several conserved structural proteins - spike (S), envelope (E), membrane (M), and nucleocapsid (N).

Coronaviruses also encode unconventional polypeptides; these include short upstream ORFs (uORFs) with possible regulatory roles, in-frame internal ORFs (iORFs) generating N-terminally truncated proteins, and out-of-frame iORFs producing novel, alternate proteins. Collectively, these ‘noncanonical’ proteins increase the coding capacity of an otherwise limited genome. One study identified 23 previously unannotated ORFs across the SARS-CoV-2 genome^3^, greatly expanding the protein repertoire of the virus. At least one noncanonical protein has been shown to have a clear functional role. The SARS-CoV-2 protein ORF9B, encoded by an out-of-frame iORF nested inside the N gene, antagonizes MAVS-mediated innate immune signaling through its interactions with TOM70^4–6^. Notably, ORF9B is conserved throughout *Betacoronaviridae*, suggesting noncanonical translation is a general feature of coronavirus genomes.

Like all viruses, coronavirus is completely dependent on the cellular translation machinery for the synthesis of viral proteins. Upon infection, SARS-CoV-2 blocks cellular protein synthesis and hijacks the translation apparatus for its own use^7–9^. At the same time, infected cells attempt to suppress bulk translation while selectively translating antiviral genes^10–12^. This tug-of-war battle plays a key role in SARS-CoV-2 infection and pathogenesis, but remains poorly characterized for other coronavirus species.

To address this gap, we comprehensively analyzed the transcriptional and translational landscape of HCoV-OC43 infection. We identified numerous noncanonical transcripts and several novel ORFs distributed across the viral genome. We also show that infected cells manage to mount a transcriptional and translational response to virus-induced ER stress, but classical antiviral pathways largely remain suppressed. At the same time, infected cells show increased translation of noncoding RNAs normally targeted by nonsense mediated decay.

## RESULTS

### Experimental design

To better understand the course of coronavirus infection and the host responses, we followed OC43 expression, together with cellular transcription and translation over a detailed time course (Fig. 1a). To this end, we infected human MRC-5 cells with OC43 at a multiplicity of infection (MOI) of 7 (Fig. 1a, S2a-b). Following a 1hr adsorption, the viral inoculum was removed, and samples were collected at regular intervals out to 30 hours post infection (hpi). Samples were harvested for RNAseq at each timepoint, and for riboseq analysis at most timepoints. In parallel, we ran mock infections using heat-inactivated virus. Virus-infected cells remained intact throughout the time course, but showed a reduced growth rate compared to mock-infected cells (Fig. S2c-d).

**Figure 1.**
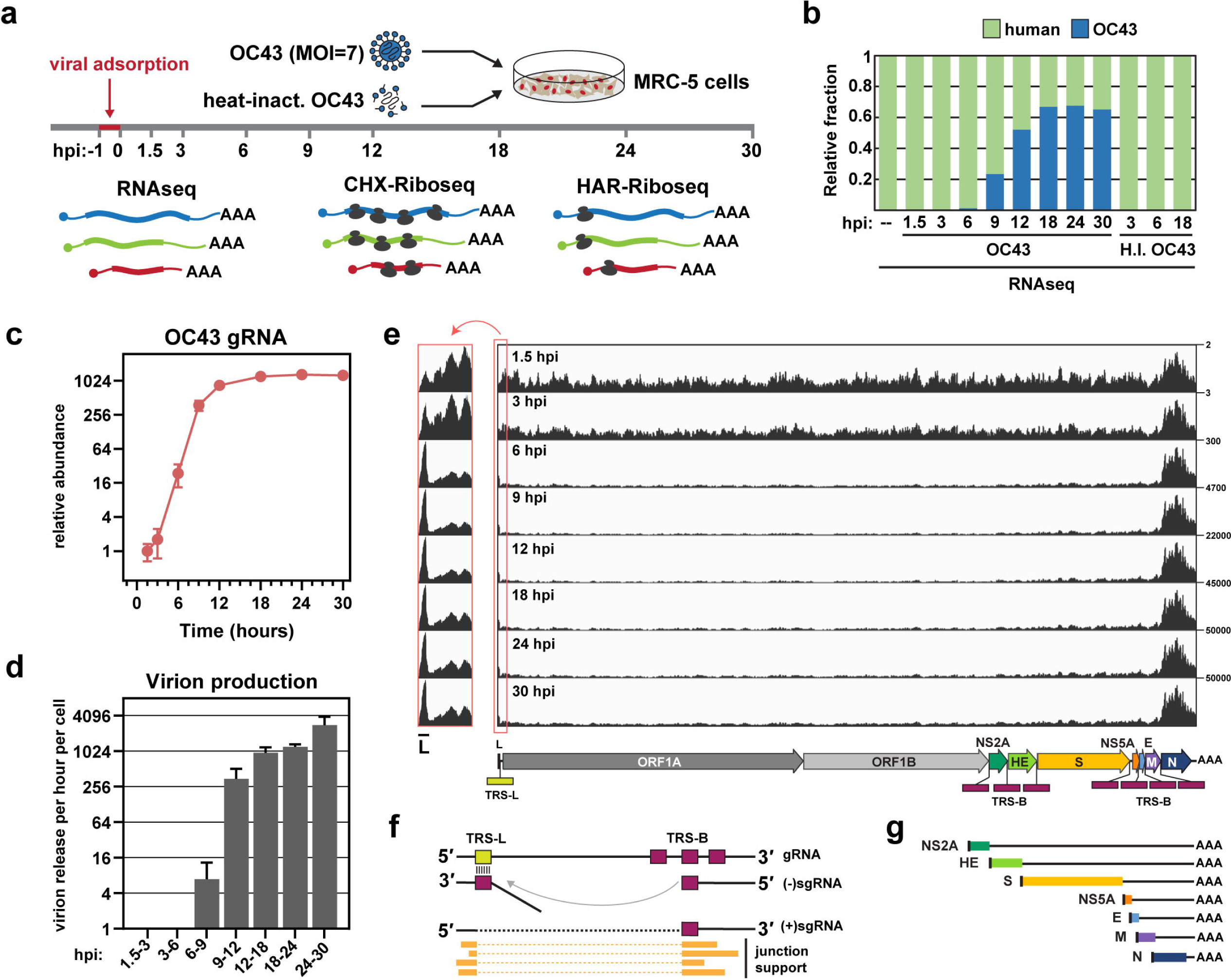
Experimental design and overview of viral transcription. (a) Overall design of the study. MRC-5 lung fibroblast cells were infected with OC43 virus at 33°C at a MOI of 7. RNA samples were collected prior to infection (-1) and at 1.5, 3, 6, 9, 12, 18, 24, and 30 hours post infection (hpi). Supernatants were also collected to assay for virion release (not shown). At most timepoints, samples were also prepared for cycloheximide (CHX)-based Riboseq. Harringtonine (HAR)-based Riboseq samples were prepared only from uninfected cells and 6 hpi. Alternatively, cells were inoculated with heat-inactivated virus, and RNA samples were analyzed at 3, 6, and 18 hpi. (b) The relative proportion of human and viral RNA throughout the course of infection. H.I. = heat inactivated. (c) Plot showing the expression kinetics of viral gRNA throughout the time course. (d) Estimated virion release per cell per hour quantified using RT-qPCR for the viral genome. Error bars show standard deviation of the mean (n=3). (e) Genome coverage tracks showing RNAseq reads across the OC43 genome. The numbers to the right indicate the scale in RPM. The structure of the full-length genomic RNA is shown below. A TRS-Body (TRS-B) is present just upstream of each of the seven ORFs in the 3′ end of the genome. The leader TRS (TRS-L) is located at the 5′ end of the genome. *Left:* The 5′ end of the genome is enlarged for clarity. (f) Coronavirus transcription/replication cycle. The template switch event is represented by the curved grey line, and the resulting junction sites are identified by sequence reads spanning the junction. (g) Schematic representation of the canonical (+)sgRNAs. The leader sequence is shown in black.

### OC43 replication and transcription kinetics

A summary of the sequencing results across the time course is presented in Fig. 1b. At later time points, viral RNA, including both genomic RNA (gRNA) and subgenomic mRNAs (sgRNAs), comprised ∼65% of all sequencing reads (Fig. 1b). By contrast, cells exposed to heat-inactivated virus showed no evidence of infection. For infected cells, intracellular levels of viral gRNA increased exponentially between 3-12 hpi, before reaching steady-state between 12-18 hpi (Fig. 1c). This plateau phase coincided with a sharp increase in viral genomes in the supernatant, suggesting that by 12-18 hpi the synthesis of new viral genomes is offset by export of mature virus particles from the cell. We measured the concentration of virions in the media using RT-qPCR, and, after accounting for the number of cells in the dish, were able to estimate virion production per cell throughout the infection cycle (Fig. 1d). Remarkably, the average cell released ∼1000 virions per hour between 12 and 24 hpi.

We next examined the distribution of sequencing reads across the viral genome (Fig. 1e). At the earliest time point (1.5 hpi; *top track*), coverage was fairly even, indicating that the viral RNA population mainly comprised full-length gRNA derived from the initial inoculum. By 6 hpi, most reads mapped to the 3′ end of the viral genome, consistent with the onset of sgRNA production. To quantify the abundance of different sgRNAs, we developed a pipeline to identify transcription junction sites. Coronavirus RdRp undergoes discontinuous transcription, in which ‘jumping’ during (-) strand sgRNA synthesis creates junctions between distant regions of the primary sequence. A second round of transcription then generates the (+)sgRNA complement. Junctions can be discriminated and quantified by sequencing and counting the number of reads spanning individual junctions across (+)sgRNAs^13^ (Fig. 1f). This approach identified seven major sgRNAs, corresponding to the seven distinct ORFs in the 3′ region of the OC43 genome (shown schematically in Fig. 1g).

Each of the seven major sgRNAs consisted of a 56-58 nt leader sequence, derived from the 5′ end of the genome, fused to a downstream open reading frame. Normalized read counts for individual sgRNAs were highly reproducible between replicates (Fig. 2a). The different sgRNAs showed similar expression kinetics and reached plateau between 9-12 hpi (Fig. 2b, S3a). As expected, N sgRNA was by far the most abundant, comprising 60-80% of all viral RNA (Fig. 2c).

**Figure 2.**
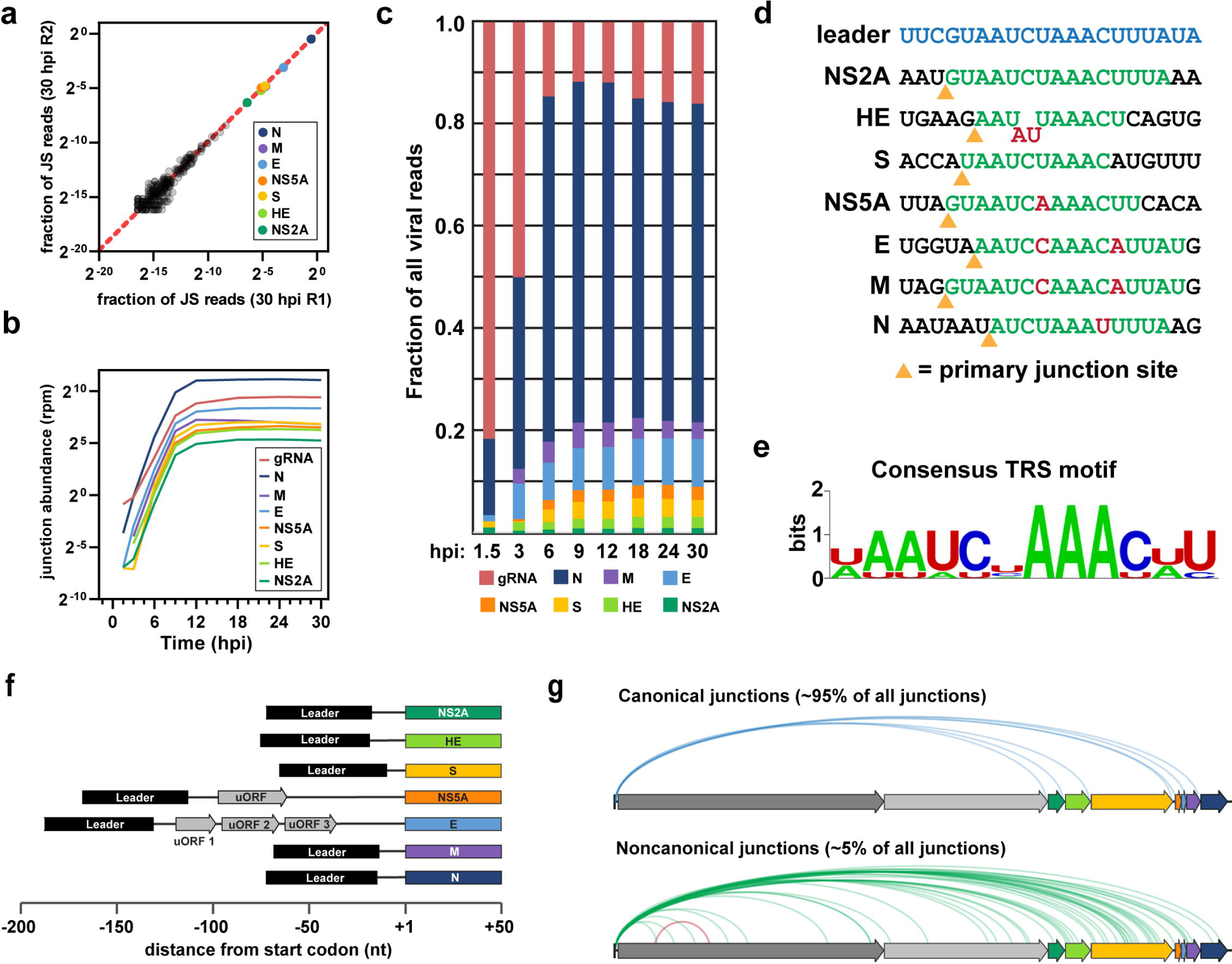
OC43 replication kinetics and subgenomic RNA expression. (a) Reproducibility of sgRNA mapping for two 30 hpi timepoints. Each dot represents the normalized read counts of a junction in replicates 1 (x-axis) and 2 (y-axis). Noncanonical junctions are shown in grey. (b) Plots showing the expression kinetics of genomic RNA and individual sgRNAs throughout the time course. Lines show the average across 3-4 replicates. (c) Histogram indicating the relative abundance of full-length genomic RNA and the seven sgRNAs at each time point. (d) Sequence context of the transcription junction site for the canonical sgRNA species. Green residues are complementary to the TRS-L, and red residues show mismatches with the TRS-L. Yellow triangles indicate the junction site. (e) The consensus TRS-B sequence. (f) Schematic representation of the 5′ UTR regions for each canonical sgRNA. (g) Sashimi plots mapping discontinuous transcription events at 30 hpi. Leader-body canonical junctions are shown in blue, leader-body noncanonical junctions are shown in green, and body-body noncanonical junctions are shown in red.

Discontinuous transcription is driven by the presence of TRS motifs, located at the 5′ end of the genome (TRS-L) and directly preceding each viral ORF (TRS-Bs) (Fig. 1e). Discontinuous transcription is thought to occur only during minus strand synthesis, when the nascent anti-TRS-B element pairs with the complementary sequence at TRS-L in the template strand, facilitating a template-switch by RdRp (Fig. S1b)^14^. For this reason, jumping efficiency at each TRS-B is partly determined by how closely it matches TRS-L^2^. Analysis of the canonical OC43 sgRNA junctions identified a consensus TRS motif of AAUCUAAAC (Fig. 2d-e), consistent with previous *in silico* predictions for OC43^15^. For five of the seven sgRNAs, the TRS-B motif was located just upstream of the start codon, resulting in relatively short 5′UTRs consisting almost entirely of the 5′ leader sequence (Fig. 2f). The NS5A and E sgRNAs were different, with junction sites much further upstream of the start codon, 114 and 132 nt respectively. The resulting 5′UTRs incorporate short uORFs which show evidence of translation (see below).

Collectively, the seven ‘canonical’ sgRNAs comprised ∼95% of all junction-spanning reads (Fig. 2g). The remaining reads were divided amongst 265 noncanonical sgRNAs, most of which lacked any substantial coding potential (Fig. S3b-c) and probably represent transcriptional noise. On average, the noncanonical transcripts had weaker donor sites (i.e. TRS-B motifs) compared to the canonical sgRNAs (Fig. S3d), consistent with their much lower accumulation. There were, however, some exceptions; for instance, the TRS-B for the HE gene was weaker than some noncanonical TRS-Bs, suggesting that additional factors contribute to jumping efficiency at particular sites. Interestingly, nearly all (99.8%) of sgRNAs used the correct acceptor site (Fig. 2g). This suggests that acceptor site selection is less error-prone than donor site selection.

### Quality control for cellular and viral Riboseq

Next, we employed ribosome profiling to characterize the OC43 translatome. We used two different translational inhibitors for our ribosome profiling experiments; cycloheximide (CHX) arrests elongating ribosomes, and harringtonine (HAR) stalls ribosomes transitioning between initiation and elongation (Fig. 3a-b). When combined with runoff of already-elongating ribosomes, HAR treatment can be used to map translation start sites^16^. In both variants, the lysate is treated with RNase I to generate ∼26-34nt mRNA fragments corresponding to the footprint of the translating ribosome, also known as ribosome protected fragments (RPFs)^17^. After library preparation, RPFs can be mapped to the human and viral genomes to identify translation initiation sites (HAR) and evaluate translation efficiency (CHX).

**Figure 3.**
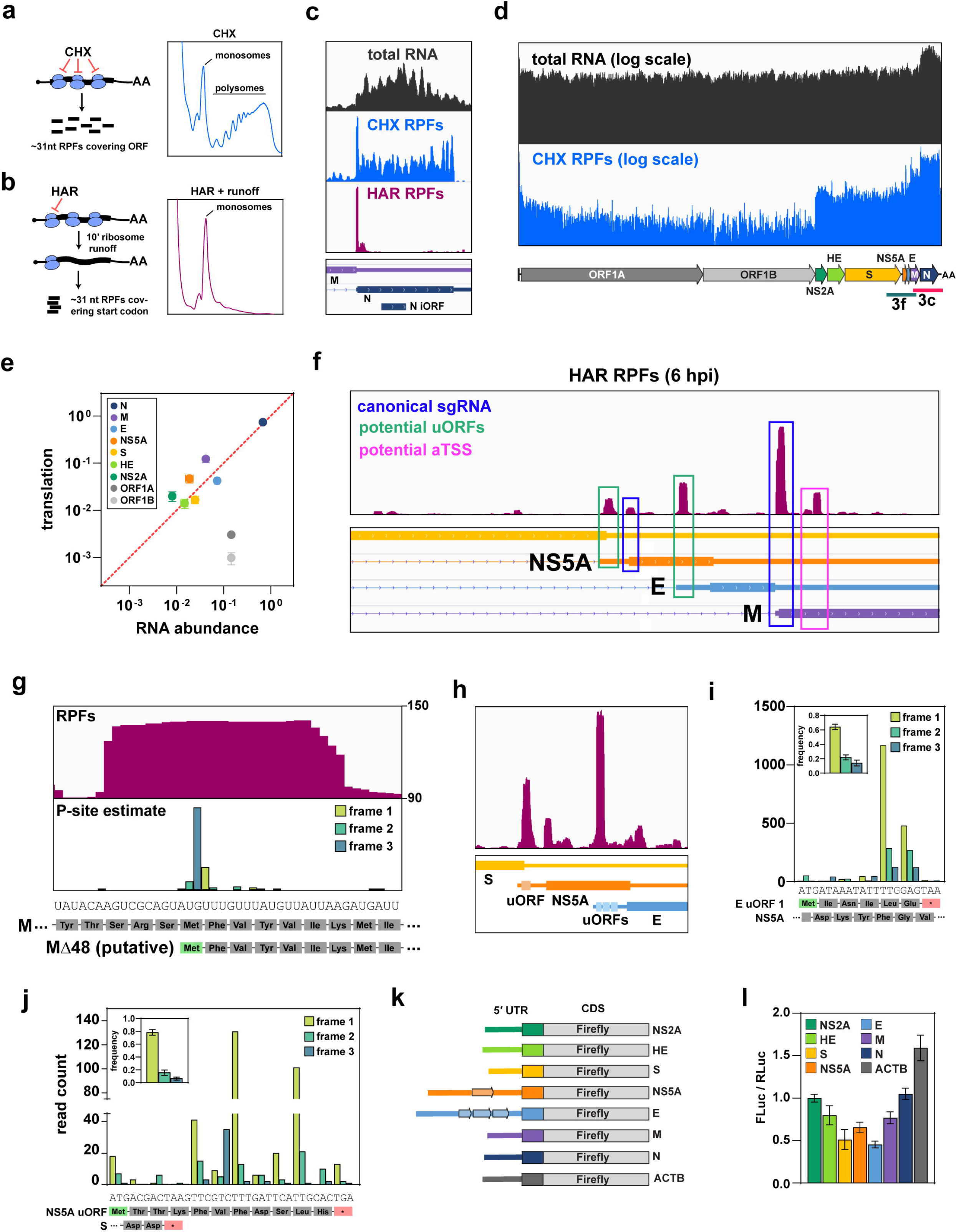
OC43 translation and protein expression. (a-b) Schematic outline of CHX- and HAR-based ribosome profiling. See text for details. *Right:* Polysome profiles following treatment with either CHX or HAR. With CHX treatment most ribosomes are found in polysomes, whereas after 10m HAR treatment to specifically block at initiation only monosomes remain. (c) Distribution of CHX RPFs (blue) and HAR RPFs (pink) across the viral N gene at 6 hpi. Total RNA (black) is included as a control. (d) A log-scale representation of CHX RPFs across the entire viral genome. The scale bars below the annotation highlight the regions displayed in (c) and (f). (e) Comparison between RNA abundance and translational output for viral genes. (f) Distribution of HAR RPFs across the sgRNA region of the genome. Prominent peaks are annotated according to their probable function. (g) A close-up view of the HAR RPF peak internal to the M gene. The inferred P-site mapping is shown below. *Below*: the codon and amino acid sequence for the full length M gene and the putative truncation variant. (h) A close-up view of the HAR RPF peaks across the NS5A and E genes. (i-j) P-site mapping for CHX RPFs across the first E uORF (i) and NS5A uORF (j). *Inset*: Summed read-frame frequencies across the entire uORF. Error bars show standard deviation of the mean (n=4). *Below*: the codon and amino acid sequence for each uORF. (k) Schematic outline of the viral 5′UTR-luciferase reporter constructs. Each construct included the indicated viral 5′UTR and first 18 nt of the corresponding viral CDS fused to firefly luciferase. (l) Bar graph showing the Firefly/Renilla (FLuc/RLuc) ratio for each construct. Error bars show the standard deviation around the mean (n=4).

Riboseq reads mapping to cellular CDS regions peaked at 31-32nt (Fig. S4a), somewhat longer than the canonical footprint size of 28nt^17^. Even so, these longer reads showed clear evidence of triplet periodicity (Fig. S4b). Inferred locations of the corresponding ribosomal P-sites aligned with the first position within the codon (‘frame 1’), indicating faithful recovery of ribosome occupancy. We also examined the distribution of reads between cellular 5′UTR, CDS, and 3′UTR regions (Fig. S4c-d). As expected, CHX riboseq reads mapped predominantly to CDS, whereas HAR riboseq reads spanned the 5′UTR and CDS. At later timepoints we saw increased coverage across 3′UTRs in CHX; for most genes the effect was modest but see below. Taken together, these observations indicate that a large proportion of reads mapped to cellular mRNAs in each dataset represent genuine RPFs.

The data for reads mapped to viral RNAs were more complex. At 6 hpi, CHX RPFs showed relatively even coverage across viral ORFs and were largely excluded from the 3′UTR (Fig. 3c, S5a-d). In addition, reads mapping to ORF1A displayed clear triplet periodicity (Fig. S5e). Unfortunately, this periodicity was lost at subsequent timepoints, and read coverage became increasingly erratic. We observed a large increase in reads mapping to ORF1B (Fig. S5b), which should be translated only at low levels, and the viral 3′UTR (Fig. S5c-d), which should not be translated at all. We conclude that many of the virus-mapping reads from 12-30 hpi are not genuine RPFs, perhaps representing packaging intermediates. We therefore excluded later timepoints from our analysis of viral translation and focused on the 6 hpi timepoint, prior to substantial virion production (Fig. 1d).

### Quantitative analysis of viral translation

A genome-wide view of viral translation at 6 hpi showed relatively low ribosome density across ORF1AB, and increased density throughout the subgenomic region (Fig. 3d), reflecting the high translation rates of subgenomic mRNAs. Surprisingly, RPF coverage over the ORF1A gene was highly uneven (Fig. S6a-b); ribosome density across the first ∼2.7kb was up to 10-fold higher than the rest of ORF1A. This may reflect either slow elongation or elevated rates of premature translation termination within this region (see Discussion).

As discussed above, some proportion of ribosomes terminate at the end of ORF1A while the remainder undergo a -1 frameshift and continue translating to the end of ORF1B. To calculate the “frameshift efficiency”, we divided the footprint density in ORF1B by the density in ORF1A (excluding the region of high ribosome density across NSP1 and NSP2). We estimate a frameshift efficiency of 56±16%, similar to values measured previously for SARS-CoV-2 and _MHV_^3,18^.

To determine the translation efficiency of viral mRNAs, we compared ribosome density to transcript abundance (Fig. 3e). For most viral ORFs, translation rate correlated well with RNA abundance. The main outliers were ORF1A and ORF1B, which showed considerably lower translation than expected based on their RNA abundance. Presumably, this is because some proportion of full-length genomic RNA is used for replication or packaging, and thus not accessible to the translational machinery.

### Noncanonical translation across the viral genome

Next, we used our HAR riboseq library to catalog translation initiation sites throughout the viral genome. As expected, most HAR peaks were centered over the start codons of canonical genes, but we also identified putative noncanonical translation start sites (Fig. 3f, S6c-d). Notably, the M gene included a prominent peak over an internal, in-frame AUG (Fig. 3f-g), suggesting an alternative translation initiation site. Translation initiation from this downstream AUG is expected to yield an N-terminal truncation variant of M lacking the first 48 amino acids (Fig. 3g). By counting the number of RPFs associated with each start site, we estimate that the Δ48M variant is translated at 21% the rate of the full-length ORF (Fig. S6e).

We also observed peaks coinciding with uORFs in the 5′UTRs of the *NS5A* and *E* sgRNAs (Fig. 3f, 3h). P-site analysis revealed evidence of periodicity (Fig. 3i-j), suggesting both uORFs are actively translated. The uORFs from *NS5A* and *E* are only 11 and 6 codons, respectively, and unlikely to produce functional peptides, but may influence translation of the downstream ORF. To test this hypothesis, we generated reporter constructs in which the 5′UTR of each viral gene was fused to Firefly luciferase (Fig. 3k). The resulting constructs were transfected into uninfected HEK293A cells alongside a Renilla luciferase control, and the reporter proteins were quantified using a dual luciferase assay. The *NS5A* and *E* reporters produced less protein than some other viral 5′UTR reporters, but the differences were modest (<2-fold) (Fig. 3l). We conclude that neither the *NS5A* uORF nor the three *E* uORFs substantially impedes translation of the downstream ORF, at least in uninfected cells.

Finally, we also observed translation at a uORF upstream of ORF1AB (Fig. S7a). This uORF is highly conserved throughout *Betacoronaviridae* (Fig. S7b-d), as previously reported^18,19^. By contrast, the *NS5A* and *E* uORFs are apparently unique to OC43 (Fig. S7e).

### ER-stress response genes are upregulated during infection

We next examined changes in cellular gene expression throughout the course of infection. RNAseq replicates collected for each timepoint showed excellent reproducibility, as assessed by Spearman correlation (Fig. S8a) and principal component analysis (PCA) (Fig. 4a). These global analyses revealed that most transcriptional changes only occur relatively late in infection, from ∼12 hpi onwards (Fig. S8a-b, 4a-b) when virus production is already high (∼1000 virions h^-1^; Fig. 1d).

**Figure 4.**
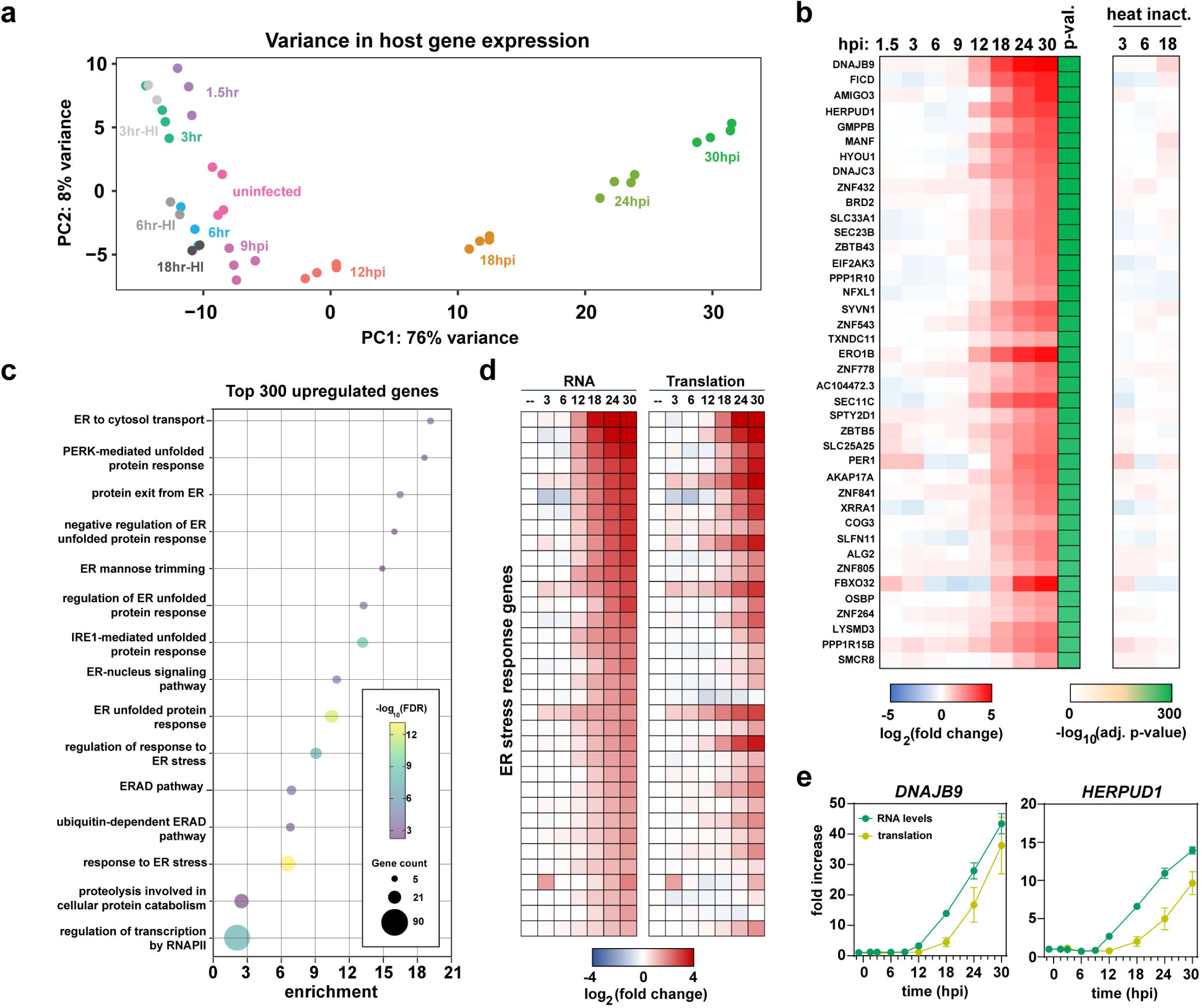
Changes to the host transcriptome and translatome during infection. (a) Principal component analysis (PCA) for different replicates and timepoints. (b) Heatmap showing the changes in RNA abundance throughout the time course. The heatmap shows the most significantly upregulated transcripts (by p-value). (c) GO enrichment analysis of the top 300 upregulated genes at 30 hpi. The size of each circle is proportional to the number of genes and the color shows the statistical significance. (d) Heatmap showing the changes in RNA abundance and translation across all genes associated with ER-related GO terms in (c). (e) Changes in mRNA abundance and translation during the infection time course for two ER stress genes. Error bars show the standard deviation around the mean (n=3-4).

To quantify changes in individual transcripts, we performed differential gene expression (DEG) analysis (Fig. S8c-d, Table S4). By 30 hpi, 14% of the transcriptome was significantly upregulated (p<0.001 and log_2_(fold change)>1) and 15% downregulated. To characterize these DEGs, we performed gene ontology (GO) enrichment analysis for the top 300 significantly upregulated genes (Fig. 4c). Of the top 15 GO terms, 13 related to the ER stress response (Fig. 4c). In general, genes associated with these categories fell into two subgroups. The first consisted of chaperones that assist in the maturation of misfolded proteins within the ER (e.g. DNAJB9, DNAJB11, PDIA4, etc.). The second included genes that regulate translation in response to ER dysfunction, including the ER-associated eIF2α kinase EIF2AK3 (PERK) and other regulators (e.g. DNAJC3, PPP1R15B). We conclude that activation of ER stress response genes is a major component of the cell’s transcriptional response to infection.

We next examined whether conventional antiviral genes are upregulated upon infection. Strikingly, neither *IFNB1* (Interferon-β) nor interferon stimulated genes (ISGs)^20^ were induced (Fig. S9a), suggesting OC43 is able to suppress interferon signaling and/or evade detection by cellular pattern recognition receptors. By contrast, several cytokine and chemokine genes were strongly activated (Fig. S9b-c). Interestingly, a similar increase was also seen after mock infection with heat-inactivated virus, but generally only at 3 and 6 hr (e.g. *CXCL3* and *IL6*). This suggests that cytokine/chemokine genes are initially induced in response to virus-laden inoculum, but sustained expression requires active infection.

Finally, we investigated whether increased transcription was accompanied by increased translation. Indeed, both ER stress response genes and cytokine/chemokine genes showed elevated translation, albeit with a ∼6hr delay compared to the transcriptional induction (Fig. 4e-f, S9b). We conclude that infected cells experience ER stress during infection and mount a transcriptional and translational response to restore homeostasis. In parallel, cells activate expression of some cytokine and chemokine genes, but interferon stimulated genes generally remain suppressed.

### Cellular NMD targets show increased translation during infection

While many of the translationally-induced genes were associated with either ER stress response or cytokine production, we also identified numerous other genes (e.g. *EIF4A2, TAF1D*, and *CCNL1*) which did not easily fit into any particular functional category (Fig. 5a, Table S6). To understand why these genes were induced upon infection, we examined the distribution of RNA and Riboseq reads across individual transcripts.

**Figure 5.**
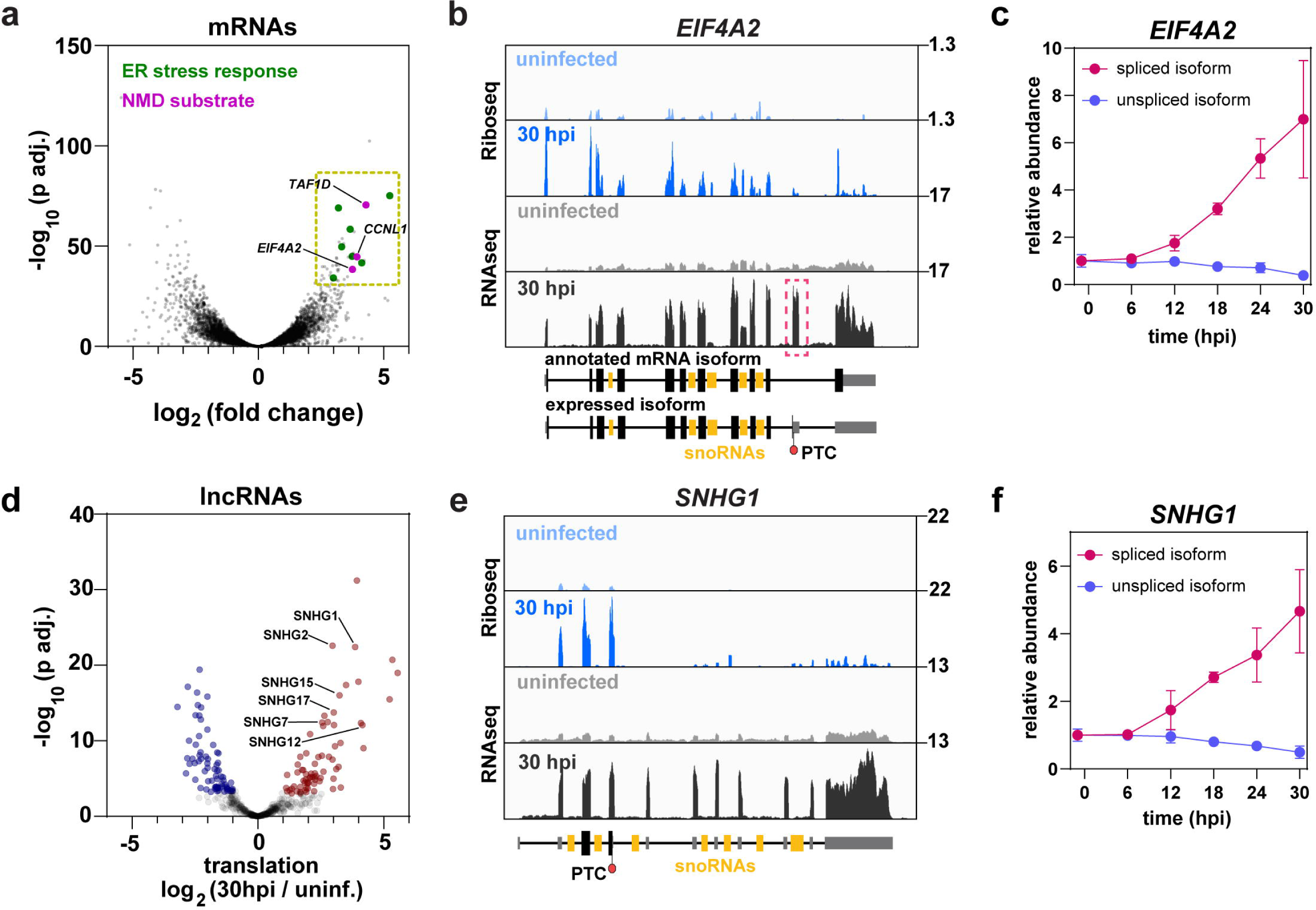
Cellular NMD targets are stabilized and translated during infection. (a) Volcano plot showing translationally upregulated (red) and downregulated (blue) mRNAs at 30 hpi. (b) Ribosome density (blue) and RNA abundance (grey) across *EIF4A2.* Numbers to the right indicate the scale in RPM. Note that *EIF4A2* incorporates an unannotated exon (dashed red box) that includes a premature termination codon (PTC). (c) Changes in RNA abundance of the spliced and unspliced isoforms of *EIF4A2*. Error bars show standard deviation around the mean (n=3-4). (d) Volcano plot showing translationally upregulated or downregulated lncRNAs at 30 hpi. (e) Ribosome density (blue) and RNA abundance (grey) across *SNHG1.* Numbers to the right indicate the scale in RPM. The location of the first termination codon is indicated as PTC. (f) Changes in RNA abundance of the spliced and unspliced isoforms of *SNGH1*. Error bars show standard deviation around the mean (n=3-4).

Intriguingly, the *EIF4A2* isoform expressed during infection includes an additional, unannotated exon with a premature termination codon (PTC) (Fig. 5b). Typically, transcripts with PTCs are recognized and degraded by nonsense mediated decay (NMD) during the first round of translation^21^. Nonetheless, *EIF4A2^PTC^* showed sharply increased translation over the course of infection. We also observed a strong increase in the abundance of spliced *EIF4A2^PTC^* RNA (Fig. 5c), presumably reflecting protection from NMD. By contrast, the unspliced precursor was either unchanged or slightly decreased. *TAF1D* and *CCNL1* followed similar patterns. The *TAF1D* isoform expressed during infection includes multiple exons downstream of the stop codon (Fig. S10a), and *CCNL1* incorporates an additional exon with a PTC (Fig. S10b), so both transcripts are predicted NMD substrates. Despite this, both *TAF1D^PTC^ and CCNL1^PTC^* showed increased translation and RNA abundance during infection. Moreover, as noted above, many genes showed increased translation across the 3’UTR (Fig. S4c), as does the viral genome (Fig. S5d) which would normally be expected to elicit an NMD response.

Next, we examined long noncoding RNAs (lncRNAs) that would also be subject to NMD if translated. We identified 76 translationally induced lncRNAs, including several snoRNA host genes (SNHGs) (Fig. 5d). SNHGs encode snoRNAs within their introns; after splicing, the mature snoRNAs are processed from the intron, while the spliced host transcript is generally degraded^22–24^. About one-half of spliced SNHG species are degraded in the nucleus by the exosome, while the remainder are exported to the cytoplasm and degraded by NMD^25^. Notably, all six of the SNHGs found here were previously identified as NMD substrates^25^. As an example, inspection of *SNHG1* revealed that translation was limited to the 5′ end of the transcript, corresponding to the first, relatively short open reading frame (Fig. 5e). The abundance of the spliced isoform of *SNHG1* was sharply increased over the course of infection, whereas the unspliced precursor was either unchanged or slightly decreased (Fig. 5f). We conclude that during OC43 infection the translation machinery becomes permissive for NMD substrates, which are translationally induced and stabilized.

## DISCUSSION

In this study, we monitored changes in RNA and translation during the course of OC43 infection. This allowed us to (1) define the transcriptional and translational architecture of the OC43 genome, and (2) characterize the cellular response to infection.

### The transcriptional and translational architecture of OC43

The OC43 transcriptome consists of a full-length genomic RNA and seven canonical sgRNAs, corresponding to the seven distinct ORFs downstream of ORF1AB (Fig. 1-2). OC43 is unusual in this regard, as related coronaviruses typically express the *NS5A* and *E* genes from a single sgRNA (Fig. S7)^18^. Beyond the seven canonical sgRNAs, we also identified 265 noncanonical sgRNAs, but these were expressed at low levels and generally lacked substantial coding potential.

Ribosome profiling allowed us to identify several well-translated novel ORFs across the viral genome (Fig. 3). Notably, the M gene apparently includes an alternative translation start site downstream of the primary AUG. Translation initiation from this distal site is expected to yield an N-terminally truncated isoform lacking the first 48 amino acids (Δ48M). We also identified several translated uORFs upstream of ORF1AB, NS5A, and E. These uORFs may produce functional peptides and/or have regulatory functions, perhaps allowing increased translation of the primary ORF when bulk translation is repressed late in infection.

To our surprise, ribosome occupancy across ORF1A was highly uneven: ribosome density at the 5′ end of the gene was at least 10-fold higher compared to downstream regions (Fig. S6). This could reflect slow elongation, perhaps caused by inefficient codons, but we failed to detect any difference in codon usage across ORF1A. An alternative explanation is that the decrease in ribosome density is caused by premature translation termination. Many viruses employ premature translation termination as a gene expression strategy, but this usually involves dedicated frameshift elements and occurs at a single, designated site. By contrast, we observed a gradual decrease in ribosome density across a relatively large region (∼2.7kb). A third possibility is that the increased ribosome density at the 5′ end reflects collisions between the viral RdRp and ribosomes.

A key feature of coronavirus replication is that the (+) strand genome serves as a template for both transcription and translation, in opposite directions. When RdRp and a ribosome both initiate on a single RNA, they will inevitably collide. We predict that collision slows or removes the leading ribosome, causing downstream depletion and/or accumulation of upstream ribosomes. We note that synthesis of the full length (-) gRNA requires transcription through the entire ORF1a/b region, which is actively translated. RdRp may require the ability to displace translating ribosomes, particularly early in infection when a single, ∼30 kb (+) gRNA molecule is the only template for both activities.

At timepoints after 6hpi, the quality of ribosome profiling data declined sharply for viral, but not cellular transcripts, probably caused by packaging of the viral genome. When treated with RNase I, partially packaged fragments will presumably yield a range of footprint sizes, overlapping with the 26-34nt fragments produced by translating ribosomes.

### Cellular response to OC43 infection

The host antiviral response was strikingly muted, with major transcriptional changes only apparent from ∼12 hpi onwards (Fig. 4). By this point, infected cells were already releasing copious quantities of virus, highlighting the extent to which OC43 is able to evade detection. Indeed, aside from several cytokine genes, the large majority of canonical antiviral genes (most notably IFNB1 and ISGs) remained quiescent throughout the entire course of infection.

In contrast, we saw activation of multiple genes linked to ER stress. We anticipate that a major, and perhaps unavoidable, consequence of OC43 infection is activation of an ER stress response. Several highly-abundant coronavirus proteins are matured through the ER, even as the virus siphons off ER membrane to build its envelope^26^. Combined, these activities may overwhelm the ER’s protein folding capacity, leading to the accumulation of misfolded proteins and triggering the unfolded protein response (UPR), a cellular pathway meant to alleviate ER stress^27^. The UPR has several downstream effects, including the upregulation of chaperone proteins and the inhibition of global protein translation to reduce the load on the ER. Beginning 12-18 hpi, ER stress response genes were both transcriptionally and translationally induced (Fig. 4). A recent study of HCoV-229E and MERS infection also reported increased transcription of ER stress response genes, but with unaltered protein expression^28^. Our results indicate that responses to OC43 are different, although we cannot exclude effects from cell lines or experimental conditions.

Finally, we observed increased translation and expression of cellular NMD substrates (Fig. 5), suggesting NMD is suppressed during infection. This may protect viral RNA from degradation, as coronavirus RNAs incorporate various features which could trigger NMD, such as multiple ORFs and exceptionally long 3′UTRs^29,30^. Indeed, the mouse coronavirus MHV was recently reported to suppress NMD *in trans*^31^. Notably, this activity is critical for viral replication, suggesting inactivation of NMD is an important and conserved feature of coronavirus infection.

## ACKNOWLEDGEMENTS

We thank Richard Clark and Angie Fawkes from the Edinburgh Wellcome Clinical Research Facility for sequencing services, Shaun Webb from the WCB bioinformatics core facility for maintaining the servers we used for processing sequencing data, and Eric Schirmer for helpful advice. This work was facilitated by the Edinburgh Protein Production Facility (EPPF). SB, ES, CD and DT were supported by Wellcome Principal Research Fellowships (109916, 222516). Work in the Wellcome Centre for Cell Biology was supported by Centre Core Grants (092076 and 203149). AA was supported by an EMBO Scientific Exchange Grant.

## DATA AVAILABILITY

The accession number for all sequence data reported in this paper is GSE252692.

## MATERIALS AND METHODS

### Cells and virus

MRC-5 cells (ATCC Cat#CCL-171) were cultured at 37°C and 5% CO_2_ in Dulbecco’s modified Eagle medium (Life Technologies Cat#41965039) supplemented with 10% fetal bovine serum (FBS; Sigma Cat#F7524) and 100U/mL penicillin and streptomycin (P/S) (Gibco Cat#15140-122). One to two days prior to infection, MRC-5 cells were transferred to 33°C. HEK293A cells (ThermoFisher Cat#R7507) were used for dual luciferase assays, and cultured in the same medium but supplemented with 1X MEM NEAA (Gibco Cat#11140-035). An aliquot of HCoV-OC43 was purchased from ATCC (Cat#VR-1558) and ∼100µL was used to infect five T-175 flasks of ∼80% confluent MRC-5 cells. Four days post infection, the medium supernatant was pooled and then centrifuged to remove cell debris. The virus-laden supernatant was divided into aliquots and frozen at -80°C. The concentration of the viral stock was calculated using qPCR (see below) and infectivity was measured by serial dilution and visualization of cytopathic effect in MRC-5 cells. For mock infections, the virus stock was heat-inactivated at 60°C for 10 min. All HCoV-OC43 virus work was conducted in a Category 2 facility in accordance with the biosafety guidelines at the University of Edinburgh.

### Plasmids

The viral 5′UTR reporter constructs were synthesized using GeneArt and cloned into pLV-SARS-CoV-2 5′UTR-Luciferase (Addgene Cat#191480)^32^ using NdeI and BsaBI. The hCMV-IE1:Renilla plasmid (Addgene Cat#118066)^33^ was used as a control for the dual luciferase assays.

### Immunofluorescence

MRC-5 cells were seeded to 4-well 0.5 ml chamber slides (Ibidi Cat#80426) and fixed at 6 hr intervals following infection. Briefly, chamber slides were washed with PBS and incubated with 4% formaldehyde for 15 min. Subsequently, cells were washed with PBS and permeabilized with 0.2% Triton X-100 for 7 min. Cells were incubated with 0.2% fish gelatin blocking solution (Biotium Cat#22010) for 30 min before addition of sheep anti-N antibody (MRC Protein Phosphorylation and Ubiquitylation Unit Cat#DA116) for 1 hr. Finally, chamber slides were washed with PBS and incubated in darkness with an anti-sheep secondary antibody conjugated to Alexa Fluor dyes (Abcam Cat#ab150177) and DAPI at a dilution of 1:5000. Chamber slides were mounted with fluoromount G (EM Sciences Cat#1798425). Images were obtained using a Zeiss Axio Imager equipped with 0.65 NA 40x objective.

### qPCR

To measure the concentration of viral genomes in the medium, we first made a plasmid incorporating a small region of ORF1AB for use as a quantification standard. This plasmid was used to generate a qPCR standard curve relating plasmid DNA concentration to the cycle threshold (Ct) value of the ORF1AB fragment. The reaction consisted of 0.5 µL Luna RT (Promega Cat#E3005S), 2.5 µL of 2X reaction mix, 300 nM of primers (Table S1) targeting ORF1AB, 1.75 µL template, and 0.5 µL water. To measure virion concentration, 20 µL of culture medium was removed from infected cells and RNA was extracted with 1mL of trizol. The RNA was resuspended in 500 µL water, and used as template for a RT-qPCR reaction as described above.

### RNAseq

For RNAseq, approximately 7×10^5^ cells growing on 6-well plates were washed with PBS and harvested with Tri Reagent (Sigma Cat#T9424). Total RNA was extracted, and 3 µg of RNA from each sample was treated with 2U Turbo DNase (ThermoFisher Cat#AM2238) for 30 min at 37°C according to the manufacturer’s protocol. The resulting RNA (300 ng) was depleted of ribosomal RNA using the NEBNext rRNA Depletion Kit (NEB Cat#E6310). rRNA-depleted RNA was purified using RNAClean XP beads (Beckman-Coulter Cat#66514), and libraries were prepared using the NEBNext rRNA Depletion Kit for Human/Mouse/Rat (NEB Cat#E6310). Adapter-ligated cDNA was amplified with 11 cycles of PCR to generate libraries ∼300 bp in length. Libraries were sequenced with a 100-cycle kit in paired-end mode.

### Ribosome profiling

Approximately 1×10^7^ cells growing on a 15 cm plate were used for each ribosome profiling experiment. For harringtonine-based profiling, the cells were treated with 8 µg/mL harringtonine (Cat#ab141941) for 10 min. Subsequently, the culture medium was removed, and cells were washed with ice-cold PBS supplemented with 0.1 mg/mL cycloheximide (CHX) (Sigma Cat#C7698). The cells were then resuspended in 1 mL of lysis buffer (10 mM Tris 7.5, 5 mM MgCl_2_, 100 mM KCl, 1% tritonX-100, 2 mM DTT, protease inhibitors (1 tablet / 10 mL) (Pierce Cat#A32955), and 0.1 mg/mL CHX), and lysed by gently passing through a 25G syringe (BD Cat#300600) four times and incubating on ice for ∼5 min. Subsequently, the cell lysates were cleared by centrifugation at 1300g/10 min/4°C and frozen at -80°C for later use. Cycloheximide-based profiling was performed as described above, but the pre-treatment with harringtonine was omitted.

Subsequently, the cell lysate was thawed, and approximately 80 µg of RNA in 1 mL volume was digested with 5 U of RNase I (Lucigen Cat#E0067-10D1) for 45 min at 25°C. Ribosomal complexes were separated by centrifugation at 38000 RPM for 2 hr at 4°C in a 14 mL 10-50% sucrose gradient with a buffer consisting of 20 mM HEPES-KOH pH 7.4, 5 mM MgCl_2_, 100 mM KCl, 2 mM DTT, and 0.1 mg/mL CHX. Centrifugations were performed using the Beckman-Coulter Optima XPN-100 ultracentrifuge with the SW40 rotor and 14mL polyallomer centrifuge tubes (Seton Scientific Cat#5031). The sucrose gradients were separated into 200 µL fractions using the Piston Gradient Fractionator (Biocomp). The fractions corresponding to the monosome peak (usually 15 in total) were combined, and Tri Reagent was used to extract RNA. Approximately 10 µg of total RNA was separated on a 15% TBE-urea gel at 200V for 65 min, and 26-34 nt footprints were excised using 26 nt and 34 nt RNA markers as a guide (Table S1)^17^. The resulting gel fragments were incubated overnight on a rotating wheel in 400 µL RNA extraction buffer (300 mM NaOAc, pH 5.2; 1 mM EDTA; and 0.25% SDS). Subsequently, the extracted RNA was precipitated with 1.5 µL GlycoBlue (Life Technologies Cat#AM9515) and 500 µL isopropanol. The RNA was pelleted by centrifugation at 20000g/15 min/4°C, washed with 80% ethanol, and resuspended in 12 µL water.

Subsequently, the RNA footprints were treated T4 PNK to generate 5⍰ and 3⍰ ends amenable to linker ligation. The 12 µL RNA was incubated together with 1.5 µL 10X buffer, 0.5 µL T4 PNK (NEB Cat#M0201), and 0.5 µL RNasIN (Promega Cat#N2511) at 37°C/30m. Next, 1.5 µL of 10 mM ATP was added and the reaction was incubated for another 30m. The RNA was purified by standard phenol:chloroform extraction, precipitated in 70% ethanol, and resuspended in 6 µL. The resulting RNA was used as input for the SMARTer smRNA-Seq Kit for Illumina (Takara Cat# 635030). cDNA libraries were prepared according to the manufacturer’s protocol, except an rRNA depletion step was incorporated after the reverse transcriptase reaction. Briefly, 12 biotinylated oligos were designed to target abundant rRNA fragments produced by RNase I cleavage (Table S1). Individual 100 µM stocks were prepared, and then combined in equivolume proportions to prepare the subtractive oligo mix. Subsequently, 20 µL cDNA was combined with 2.5 µL subtractive oligo mix and 2.5 µL 20X SSC (3 M NaCl, 0.3 M NaCitrate, pH 7). The resulting reaction was denatured at 100°C for 90s, and then cooled to 37°C at a rate of 0.1°C/s to allow annealing. Subsequently, the reaction mix was combined with 50 µL MyOne Streptavidin C1 Dynabeads (Life Technologies Cat#65001) in 2X B&W buffer (2 M NaCl, 1 mM EDTA, 10 mM Tris 7.5). The resulting reaction was incubated at 37°C for 15m with shaking. The dynabeads were pelleted using a magnetic rack and 40 µL of the supernatant was recovered and transferred to a new tube containing 1.5 µL GlycoBlue, 6 µL of 5 M NaCl, and 60 µL water. The resulting mix was combined with 150 µL isopropanol, and incubated on ice for 30 min. The rRNA-depleted cDNA was pelleted at 20000g/30 min/4°C, washed with 80% ethanol, and resuspended in 20 µL water.

Finally, 5 µL was used as input for a 13-cycle PCR reaction according to the manufacturer’s protocol. The resulting PCR product was purified using the Nucleospin Gel and PCR clean-up kit (Macherey-Nagel Cat#740609.50), and resolved on a 8% polyacrylamide TBE gel (Life Technologies Cat#EC62162BOX). Fragments from ∼190-250bp in length were excised, and incubated overnight on a rotating wheel in 400 µL DNA extraction buffer (300 mM NaCl, 10 mM Tris 8.0, 1 mM EDTA, and 0.01% igepal). Subsequently, the extracted DNA was precipitated as described above for the RNA footprints. Libraries were sequenced using Illumina NextSeq with a 100-cycle kit in single-end mode.

### Reporter assays

HEK293A cells were seeded onto 24 well plates and transfected with individual 5′UTR reporters together with the Renilla luciferase control the following day. Plasmids were combined in a 1:1 ratio (250 ng each) and transfected using Lipofectamine 2000 according to the manufacturer’s protocol. The following day, cells were harvested using 200 µL Passive Lysis buffer. Firefly and Renilla luciferase production were evaluated using the Dual Luciferase Reporter Assay System (Promega Cat#E1910) and the Multimode Plate Reader (Molecular Devices).

## QUANTIFICATION AND STATISTICAL ANALYSIS

### RNAseq analysis

Sequencing reads were aligned to a concatenated human (ENSEMBL hg38-101) and HCoV-OC43 (ATCC_VR_1558) genome using `STAR`^34^. The resulting bam files were sorted (`samtools sort`) and used to generate coverage maps with `bamCoverage`. The coverage maps were normalized based on cellular mRNA content, so the viral genome was excluded from the quantification:

`bamCoverage -b file.bam --ignoreForNormalization HCoV_OC43 -- exactScaling --normalizeUsing CPM --binSize 1 --filterRNAstrand forward -o file_plus_strand.bw`

Read counts per gene were tabulated using featureCounts^35^:

`featureCounts file.bam -a human_OC43_annotation.gtf -C -d 50 -D 2000 - -countReadPairs -s 2 -p -B -O --extraAttributes gene_biotype,gene_name --largestOverlap -t gene -o read_counts.txt`

Virus junction counts were based on the number of uniquely mapped reads spanning discontinuous regions of the viral genome. These were identified by extracting reads with gapped alignments from the sam file.

### Riboseq analysis

The Takara SMARTer smRNA-Seq Kit adds 3 nt to the 5⍰ end of each read, and a short poly(A) tail at the 3⍰ end. These features were removed using flexbar:

`flexbar -r file.fastq -n 10 -a AAAAAAAAAA -ao 4 -x 3 -m 1 -ae RIGHT -u 2 -f i1.8 -t filtered.fastq`

The remaining reads were aligned to the concatenated human and HCoV-OC43 genome as described above. The coverage maps were normalized based on cellular mRNA content, so the viral genome and chromosomes encoding ribosomal RNA (Chr 1, 21, and 22) were excluded from the quantification:

`bamCoverage -b file.bam --ignoreForNormalization 1 21 22 HCoV_OC43 -- exactScaling --normalizeUsing CPM --binSize 1 --filterRNAstrand forward -o file_plus_strand.bw`

Read counting was performed with featureCounts:

`featureCounts file.bam -a human_OC43_annotation.gtf -C -d 50 -D 2000 - -countReadPairs -s 1 -O --extraAttributes gene_biotype,gene_name -- largestOverlap -t gene -o read_counts.txt`

Basic quality control analyses for cellular mRNAs (Fig. S4-5; RPF lengths, frame, and biotypes) were performed with Ribotoolkit^36^ (http://rnainformatics.org.cn/RiboToolkit/index.php) using default parameters. For frame and biotype analysis, only 26-34nt reads were analyzed.

Viral mRNAs were analyzed with Ribocount (https://pythonhosted.org/riboplot/ribocount.html) using a fasta annotation file corresponding to the viral ORFs. A 12 nt offset was used for all RPFs 26-34 nt in length:

`ribocount -b file.bam -f OC43_CDS.fasta -l 26,27,28,29,30,31,32,33,34 -s 12,12,12,12,12,12,12,12,12 -o OC43_RPFs.txt`

### Differential gene expression

DESeq2^37^ was used to identify differentially expressed genes and produce PCA and volcano plots. As input, we used the top 14000 cellular mRNAs by expression (defined as the average RPM across all timepoints and replicates). Default parameters were used for all analyses. DEGs were defined as having a log_2_(fold change)>±1 and (adj. p-value)<0.001.

## FIGURE LEGENDS

**Figure S1. Related to Introduction.** (a) The viral life cycle. See Introduction for details. (b) Schematic outline of discontinuous transcription. The viral genome incorporates TRS-B motifs upstream of each structural and accessory protein ORF. When RdRp transcribes a TRS-B, it sometimes “jumps” to the TRS-L at the 5′ end of the genome. Because the TRS-L and each TRS-B are similar in sequence, the nascent RNA reanneals to the template, and transcription resumes. The relative abundance of each (-) sgRNA is determined by how frequently RdRp jumps at each TRS-B. In the final step, the (-) sgRNAs serve as templates for synthesis of (+) sgRNAs which are subsequently used in translation.

**Figure S2. Experimental design and validation. Related to** Figure 1. (a-b) Preparation of the viral stock. To minimize experimental variability, we prepared a single batch of virus for the duration of the study. (a) Viral genomes were quantified using RT-qPCR against a region within ORF1A. (b) Infectious particles were quantified using limiting dilution and assessment of cytopathic effect (CPE) under the experimental conditions used in the study. The resulting particle:infection ratio was 7:1. (c) Immunofluorescence microscopy showing infected cells throughout the time course. Cells were stained with DAPI to visualize cell nuclei and decorated with an antibody against the viral N protein to identify infected cells. (d) Cell growth rates following infection with OC43 or mock-infection with heat-inactivated (h.i.) virus. Error bars show the standard deviation around the mean (n=3).

**Figure S3. OC43 noncanonical sgRNA expression. Related to** Figure 2. (a) Plots showing the expression kinetics of genomic RNA and individual sgRNAs throughout the time course. Error bars show standard deviation of the mean (n=4). (b) Tree diagram showing the structure of the 50 most abundant noncanonical sgRNAs. (c) Structure and protein coding potential of the eight most abundant noncanonical sgRNAs. In every case, the initial ORF is no longer than 16 amino acids. The numbers indicate the donor-acceptor coordinates. (d) Plot showing the hybridization potential between donor sites (i.e. TRS-Bs) and the TRS-L acceptor site. Transcripts are sorted by abundance. Complementary bases are displayed in orange (canonical junctions) or blue (noncanonical junctions), and noncomplementary bases are shown in white.

**Figure S4. Riboseq quality control metrics for cellular genes. Related to** Figure 3. (a) Length distribution of RPFs mapping to cellular (green) and viral (blue) CDS regions following gel-based selection for 26-34 nt fragments. (b) Proportion of cellular-mapping RPFs associated with each reading frame. (c) Distribution of Riboseq reads between 5′UTR, CDS, and 3′UTR regions. (d) Riboseq read densities across the *EEF2* gene for a representative replicate. Total RNA (black), CHX RPFs (blue), HAR RPFs (pink). The numbers to the right of the plot show the scale in RPM for each track. The annotation for *EEF2* is shown below. Grey rectangles show 5′ and 3′UTR regions, black rectangles show CDS, and black lines show introns.

**Figure S5. Riboseq quality control metrics for viral genes. Related to** Figure 3. (a) Schematic showing the structure of the full-length viral gRNA. The scale bars below the annotation highlight the regions displayed in panels B and C. (b-c) Ribosome density across sections of the viral genome. (d) The relative proportion of viral-mapping RPFs which map to the 3′UTR. Error bars show standard deviation of the mean (n=3). (e) Proportion of viral-mapping RPFs associated with each reading frame across ORF1A. Error bars show standard deviation of the mean (n=3).

**Figure S6. OC43 translation. Related to** Figure 3. (a) Ribosome density at 6 hpi across a section of ORF1A. Four independent replicates of CHX Riboseq are shown (blue). Total RNA determined by RNAseq is shown as a comparison (black). Codon optimality scores are shown below (green). (b) Ribosome density across ORF1A. The red baseline shows the average ribosome density across the latter half of ORF1A. (c) Distribution of HAR RPFs around the start codons of each canonical viral ORF. (d) Density of HAR RPFs at 6 hpi across the subgenomic region of the viral genome. Two independent replicates are shown. The asterisk marks a likely sequencing artifact. (e) Bar chart showing the relative proportion of RPFs associated with the primary start codon for M versus the downstream initiation site. Error bars show the standard deviation around the mean (n=2).

**Figure S7. Viral uORF translation. Related to** Figure 3. (a) P-site mapping for CHX RPFs across the ORF1AB uORF. *Below*: the codon and amino acid sequence for the ORF1AB uORF. (b-d) Schematic showing the conserved architecture (b), peptide sequences (c), and sequence context (d) of the ORF1AB uORF across the betacoronavirus A lineage. (e) Genomic organization of the NS5A and E genes across various representatives of the betacoronavirus A lineage. The architectures for MHV^18^ and OC43 were determined experimentally, and HKU24 and HKU1 were inferred based on the presence or absence or TRS-B motifs.

**Figure S8. Changes to the host transcriptome during infection. Related to** Figure 4. (a) Matrix showing the pairwise Spearman correlation coefficients for human transcripts between different replicates and timepoints. The red squares outline each set of replicates. (b) Heatmap showing the changes in RNA abundance throughout the time course. The heatmap shows the most significantly decreased transcripts (by p-value). (c) Bar chart showing the number of differentially expressed genes (DEGs) at each timepoint. (d) Volcano plots showing statistically significantly upregulated (red) and downregulated (blue) genes at various timepoints after infection.

**Figure S9. Antiviral gene expression during infection. Related to** Figure 4. (a) Box-and-whisker plot showing the expression of interferon stimulated genes (ISGs) in response to infection. The box shows the 25^th^, median, and 75^th^ percentile, while the whiskers show the 10^th^ and 90^th^ percentiles. (b) Transcriptional (*left*) and translational (*center*) induction of cytokines and chemokines in response to infection. The transcriptional response to mock infection with heat-inactivated (HI) virus is also shown (*right*). (c) Changes in mRNA abundance and translation during the infection time course for two cytokine genes. Error bars show the standard deviation around the mean (n=3-4).

**Figure S10. NMD substrates are translationally derepressed during infection. Related to** Figure 5. (a-b) Ribosome density (blue) and RNA abundance (grey) across *TAF1D* (a) and *CCNL1* (b). Numbers to the right indicate the scale in RPM. Note that *TAF1D* includes eight additional exons beyond the normal stop codon, while *CCNL1* incorporates an unannotated exon that includes a PTC. Both features are highlighted with a dashed red box.

## REFERENCES

1. Forni, D., Cagliani, R., Clerici, M. & Sironi, M. Molecular Evolution of Human Coronavirus Genomes. Trends Microbiol 25, 35–48 (2017).

2. Hartenian, E. et al. The molecular virology of coronaviruses. J Biol Chem 295, 12910–12934 (2020).

3. Finkel, Y. et al. The coding capacity of SARS-CoV-2. Nature 589, 125–130 (2021).

4. Gao, X. et al. Crystal structure of SARS-CoV-2 Orf9b in complex with human TOM70 suggests unusual virus-host interactions. Nat Commun 12, 2843 (2021).

5. Thorne, L.G. et al. Evolution of enhanced innate immune evasion by SARS-CoV-2. Nature 602, 487–495 (2022).

6. Jiang, H.W. et al. SARS-CoV-2 Orf9b suppresses type I interferon responses by targeting TOM70. Cell Mol Immunol 17, 998–1000 (2020).

7. Finkel, Y. et al. SARS-CoV-2 uses a multipronged strategy to impede host protein synthesis. Nature 594, 240–245 (2021).

8. Thoms, M. et al. Structural basis for translational shutdown and immune evasion by the Nsp1 protein of SARS-CoV-2. Science 369, 1249–1255 (2020).

9. Schubert, K. et al. SARS-CoV-2 Nsp1 binds the ribosomal mRNA channel to inhibit translation. Nat Struct Mol Biol 27, 959–966 (2020).

10. Li, Y. et al. SARS-CoV-2 induces double-stranded RNA-mediated innate immune responses in respiratory epithelial-derived cells and cardiomyocytes. Proc Natl Acad Sci U S A 118(2021).

11. Aloise, C. et al. SARS-CoV-2 nucleocapsid protein inhibits the PKR-mediated integrated stress response through RNA-binding domain N2b. PLoS Pathog 19, e1011582 (2023).

12. Zheng, Z.Q., Wang, S.Y., Xu, Z.S., Fu, Y.Z. & Wang, Y.Y. SARS-CoV-2 nucleocapsid protein impairs stress granule formation to promote viral replication. Cell Discov 7, 38 (2021).

13. Kim, D. et al. The Architecture of SARS-CoV-2 Transcriptome. Cell 181, 914–921 e10 (2020).

14. Sola, I., Almazan, F., Zuniga, S. & Enjuanes, L. Continuous and Discontinuous RNA Synthesis in Coronaviruses. Annu Rev Virol 2, 265–88 (2015).

15. Lau, S.K. et al. Molecular epidemiology of human coronavirus OC43 reveals evolution of different genotypes over time and recent emergence of a novel genotype due to natural recombination. J Virol 85, 11325–37 (2011).

16. Ingolia, N.T., Brar, G.A., Rouskin, S., McGeachy, A.M. & Weissman, J.S. The ribosome profiling strategy for monitoring translation in vivo by deep sequencing of ribosome-protected mRNA fragments. Nat Protoc 7, 1534–50 (2012).

17. McGlincy, N.J. & Ingolia, N.T. Transcriptome-wide measurement of translation by ribosome profiling. Methods 126, 112–129 (2017).

18. Irigoyen, N. et al. High-Resolution Analysis of Coronavirus Gene Expression by RNA Sequencing and Ribosome Profiling. PLoS Pathog 12, e1005473 (2016).

19. Wu, H.Y., Guan, B.J., Su, Y.P., Fan, Y.H. & Brian, D.A. Reselection of a genomic upstream open reading frame in mouse hepatitis coronavirus 5’-untranslated-region mutants. J Virol 88, 846–58 (2014).

20. Schoggins, J.W. et al. A diverse range of gene products are effectors of the type I interferon antiviral response. Nature 472, 481–5 (2011).

21. Schweingruber, C., Rufener, S.C., Zund, D., Yamashita, A. & Muhlemann, O. Nonsense-mediated mRNA decay - mechanisms of substrate mRNA recognition and degradation in mammalian cells. Biochim Biophys Acta 1829, 612–23 (2013).

22. Kiss, T. & Filipowicz, W. Exonucleolytic processing of small nucleolar RNAs from pre-mRNA introns. Genes Dev 9, 1411–24 (1995).

23. Tycowski, K.T., Shu, M.D. & Steitz, J.A. A mammalian gene with introns instead of exons generating stable RNA products. Nature 379, 464–6 (1996).

24. Lykke-Andersen, S. et al. Human nonsense-mediated RNA decay initiates widely by endonucleolysis and targets snoRNA host genes. Genes Dev 28, 2498–517 (2014).

25. Bresson, S.M., Hunter, O.V., Hunter, A.C. & Conrad, N.K. Canonical Poly(A) Polymerase Activity Promotes the Decay of a Wide Variety of Mammalian Nuclear RNAs. PLoS Genet 11, e1005610 (2015).

26. V’Kovski, P., Kratzel, A., Steiner, S., Stalder, H. & Thiel, V. Coronavirus biology and replication: implications for SARS-CoV-2. Nat Rev Microbiol 19, 155–170 (2021).

27. Hetz, C. & Papa, F.R. The Unfolded Protein Response and Cell Fate Control. Mol Cell 69, 169–181 (2018).

28. Shaban, M.S. et al. Multi-level inhibition of coronavirus replication by chemical ER stress. Nat Commun 12, 5536 (2021).

29. Buhler, M., Steiner, S., Mohn, F., Paillusson, A. & Muhlemann, O. EJC-independent degradation of nonsense immunoglobulin-mu mRNA depends on 3’ UTR length. Nat Struct Mol Biol 13, 462–4 (2006).

30. Hurt, J.A., Robertson, A.D. & Burge, C.B. Global analyses of UPF1 binding and function reveal expanded scope of nonsense-mediated mRNA decay. Genome Res 23, 1636–50 (2013).

31. Wada, M., Lokugamage, K.G., Nakagawa, K., Narayanan, K. & Makino, S. Interplay between coronavirus, a cytoplasmic RNA virus, and nonsense-mediated mRNA decay pathway. Proc Natl Acad Sci U S A 115, E10157–E10166 (2018).

32. Vora, S.M., et al. Targeting stem-loop 1 of the SARS-CoV-2 5’ UTR to suppress viral translation and Nsp1 evasion. Proc Natl Acad Sci U S A 119(2022).

33. Sarrion-Perdigones, A. et al. Examining multiple cellular pathways at once using multiplex hextuple luciferase assaying. Nat Commun 10, 5710 (2019).

34. Dobin, A. et al. STAR: ultrafast universal RNA-seq aligner. Bioinformatics 29, 15–21 (2013).

35. Liao, Y., Smyth, G.K. & Shi, W. featureCounts: an efficient general purpose program for assigning sequence reads to genomic features. Bioinformatics 30, 923–30 (2014).

36. Liu, Q., Shvarts, T., Sliz, P. & Gregory, R.I. RiboToolkit: an integrated platform for analysis and annotation of ribosome profiling data to decode mRNA translation at codon resolution. Nucleic Acids Res 48, W218–W229 (2020).

37. Love, M.I., Huber, W. & Anders, S. Moderated estimation of fold change and dispersion for RNA-seq data with DESeq2. Genome Biol 15, 550 (2014).

